# COBRAme: A Computational Framework for Genome-Scale Models of Metabolism and Gene Expression

**DOI:** 10.1101/106559

**Authors:** Colton J. Lloyd, Ali Ebrahim, Laurence Yang, Zachary King, Edward Catoiu, Edward J. O’Brien, Joanne K. Liu, Bernhard O. Palsson

## Abstract

Genome-scale models of metabolism and macromolecular expression (ME-models) explicitly compute the optimal proteome composition of a growing cell. ME-models expand upon the well-established genome-scale models of metabolism (M-models), and they enable new and exciting insights that are fundamental to understanding the basis of cellular growth. ME-models have increased predictive capabilities and accuracy due to their inclusion of the biosynthetic costs for the machinery of life, but they come with a significant increase in model size and complexity. This challenge results in models which are both difficult to compute and challenging to understand conceptually. As a result, ME-models exist for only two organisms (*Escherichia coli* and *Thermotoga maritima*) and are still used by relatively few researchers. To address these challenges, we have developed a new software framework called COBRAme for building and simulating ME-models. It is coded in Python and built on COBRApy, a popular platform for using M-models. COBRAme streamlines computation and analysis of ME-models. It provides tools to simplify constructing and editing ME-models to enable ME-model reconstructions for new organisms. We used COBRAme to reconstruct a condensed *E. coli* ME-model called *i*JL1678b-ME. This reformulated model gives virtually identical solutions to previous *E. coli* ME-models while using ¼ the number of free variables and solving in less than 10 minutes, a marked improvement over the 6 hour solve time of previous ME-model formulations. This manuscript outlines the architecture of COBRAme and demonstrates how ME-models can be reconstructed and edited most efficiently using the software.

## Introduction

Genome-scale metabolic models (M-models) have shown significant success predicting various aspects of cellular metabolism by integrating all of the experimentally determined metabolic reactions taking place in an organism of interest [1–4]. These predictions are enabled based on the stoichiometric constraints of the organism’s metabolic reaction network and metabolic interactions with the environment. M-models are capable of accurately predicting the metabolic capabilities of an organism, but they require defined substrate input constraints and empirical metabolite measurements to make predictions of its growth capabilities. Therefore, a focus of development in the field of genome-scale models has been to increase the scope and capabilities of M-models [5].

Recently, M-models have been extended to include the synthesis of the gene expression machinery which can be used to compute the entire metabolic and gene expression proteome [6–9]. These ME-models integrate <u>M</u>etabolism and <u>E</u>xpression on the genome scale (**Figure 1**), and they are capable of explicitly computing a large percentage (> 80% in some cases) of the proteome by mass in enterobacteria [10]. In other words, not only do ME-models compute optimal metabolic flux states, as do M-models, but they also compute the optimal proteome usage required to sustain the metabolic phenotype. ME-models enable a wide range of new biological questions to be investigated including direct calculations of proteome allocation [11] to cellular processes, temperature dependent activity of the chaperone network[12], metabolic pathway usage and the effects of membrane and volume constraints [7]. Furthermore, their ability to compute the optimal proteome abundances for a given condition make them ideal for mechanistically integrating transcriptomics and proteomics data.

**Figure 1:**
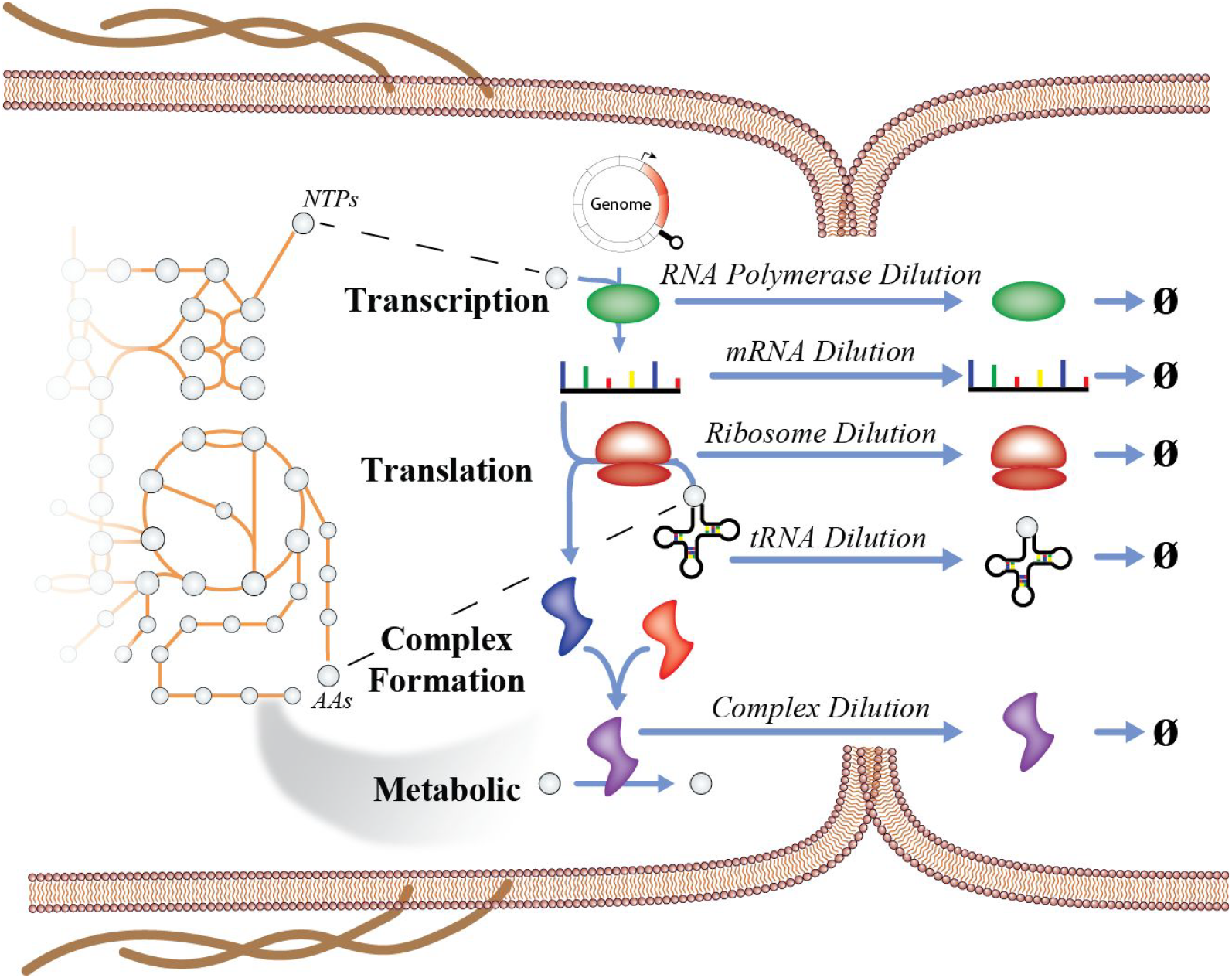
Multi-scale processes modeled in a ME-model depicted in a dividing *E. coli* cell. ME-models expand upon underlying M-models by explicitly accounting for the reactions involved in gene expression required to catalyze enzymatic processes. The synthesis of each major macromolecule is coupled to the reaction that it is involved in by accounting for its dilution to daughter cells during cell division. Each dilution reaction is a function of growth rate (μ).

So far ME-models have been constructed for only two organisms, *Thermotoga maritima* [8] and *Escherichia coli* K-12 MG1655 [6,7,9,13]. The slow pace of ME-model construction can be attributed to two basic challenges with ME-models. First, ME-models are much slower to numerically solve than M-models; it takes 5 orders of magnitude more CPU time to solve *i*OL1650-ME [6] than it does the corresponding *i*JO1366 M-model [14]. As a result, while M-models can be solved on personal computers, ME-models currently require large clusters or supercomputers to parallelize simulations. While increased computing power is generally becoming more readily available, thus alleviating the computational challenge, other challenges that come with ME-models are not as easily addressed. M-models can use generalized software tools [15–19], but each organism’s ME-model has required its own dedicated codebase and database schema, which makes advances for one organism’s model difficult to apply to another organism. Second, the large model sizes and complex structure have made analyzing and debugging the model difficult and time consuming. Therefore, each organism’s ME-model has required dedicated person-years of effort.

We addressed the above challenges by developing a computational framework written in Python – analogous to the widely used software for M-models, COBRApy [18] – for building, editing, simulating and interpreting ME-model results, called COBRAme. COBRAme is designed to: 1) be applicable to any organism with an existing metabolic reconstruction (M-model) 2) use protocols and commands familiar to current users of COBRApy 3) construct ME-models with a reaction organization that is easily interpreted by the user 4) construct models that solve orders of magnitude faster than previous ME-models [6]. As a result of the above considerations, we hope that COBRAme and its associated tools, presented here, will accelerate the development and use of models of metabolism and expression.

## Design and Implementation

### Python

The COBRAme software is written entirely in Python and requires the COBRApy [18] software package to enable full COBRA model functionality. Additionally, COBRAme requires the SymPy Python module [20] in order to handle “μ”, the symbolic variable representing cellular growth rate, which participates as a member of many stoichiometric coefficients in the ME-matrix. The BioPython package [21] is used by COBRAme to construct transcription, translation and tRNA charging reactions for each gene product in the organism’s genbank genome annotation file. The ME-model is solved using the SoPlex [22,23] or quadMINOS (Ma et al. 2017) solvers via an API written in Python and included as part of this project. Further, the ECOLIme Python package is included in this work and contains information pertaining to *E. coli* gene expression and scripts to build *i*JL1678b-ME starting with the *E. coli* metabolic model, *i*JO1366 [14]. ECOLIme can further act as a blueprint for ME-model reconstructions of new organisms.

### ME-model Architecture

Constructing a ME-model requires assembling information pertaining to many different cellular processes. For instance, in order to construct a translation reaction for the ME-model, the sequence of the gene, the codon table for the organism, the tRNAs for each codon, ribosome translation rates, elongation factor usage, etc. must be incorporated. Further, several processes in the ME-model recur for many genes that are transcribed or translated due to their template-like nature [13]. To address these challenges, the COBRAme ME-model was structured to compartmentalize information for individual cellular processes. A key component of this approach was the separation of the ME-model into two major Python class types: the information storage vessels called **ProcessData** and the functional model reactions called **MEReaction**, which is analogous to the COBRApy Reaction.

#### ProcessData

COBRAme constructs ME-models that are composed of two major Python classes. The first of these is the ProcessData class, which is used to store information associated with a cellular process. The type of information contained in each ProcessData type is summarized in the **COBRAme Documentation**. This method of information storage has several advantages over alternatives such as establishing a database to query information as it is needed, which was the approach used to build previous ME-model versions. For example, this method simplifies the dissemination of the information used to construct a ME-model given that the information can now be included as part of a published ME-model without requiring the user to install and populate a database. Further, this gives the ability to compartmentalize the information based on which cellular processes it elucidates. By storing this information in Python objects, methods can be implemented to further allow data contained in each ProcessData instance to be manipulated. This method also reduces error by enabling many features to be computed using defined inputs in a consistent way. For example, the amino acid sequence for a protein can be dynamically computed and used to construct a TranslationReaction instance using a gene’s nucleotide sequence and codon table.

#### MEReaction

ME-models are multiscale in nature meaning they contain a variety of different types of reactions that operate on different timescales and have different cellular functions. Reactions therefore must be coupled together to dictate the required activity of one reaction needed to facilitate the reactions which it participates. This is done by deriving a coupling coefficient to determine the amount of a macromolecule needed to catalyze particular reactions. To facilitate this coupling and to handle the unique characteristics of each major reaction type found in cell biology, the MEReaction Python class is used. This class defines the functional ME-model reaction that inherits from the COBRApy Reaction and thus ultimately makes up the ME-matrix. In addition to the functionality of COBRApy Reactions, MEReactions contain functions to read and process the information contained in ProcessData objects and to update this information into a complete, functional reaction. Part of compiling a functional reaction also includes imposing the appropriate coupling constraints in many cases (coupling constraints detailed in **COBRAme Documentation**). These coupling constraints are imposed directly as part of the MEReaction’s update method and varies depending on the reaction type. Since MEReactions are associated with the information used to construct them through ProcessData, this codebase has the ability to easily query, edit and update the information used to construct the reaction into the MEReaction and therefore model.

#### ME-model Construction Workflow

ME-models of *E. coli* are reconstructed using the two Python packages presented here, COBRAme and ECOLIme. COBRAme contains the class definitions and necessary methods to facilitate building and editing a working ME-model. COBRAme is written to be organism agnostic such that it can be applied to ME-models for any organism. ECOLIme contains the *E. coli* specific information (e.g. the *E. coli* ribosome composition) as well as functions required to process files containing *E. coli* reaction information (e.g. the text file containing transcription unit definitions) and associate them with the ME-model being constructed. Therefore, ECOLIme is required to assemble the reaction and gene expression information that comprises *i*JL1678b-ME, and COBRAme, on the other hand, supplies the computational framework underlying the ME-model. The composition along with further demonstrations of the utility of each of these packages is outlined in the **COBRAme Documentation**.

The process of building *i*JL1678b-ME using COBRAme and ECOLIme is presented in the building script, ‘**build_me_model**’(**Figure 2**). This script goes through each of the major gene expression processes modeled in *i*JL1678b-ME and uses the functionality contained within ECOLIme to read and process all relevant information. Once the information is loaded, it is used to create and populate ProcessData instances associated with the information. Each of ProcessData instances are then linked to the appropriate MEReaction instance and updated to form a functioning ME-model (**Figure 2**). A description of the ‘**build_me_model**’ script can be found in the **COBRAme Documentation**.

**Figure 2:**
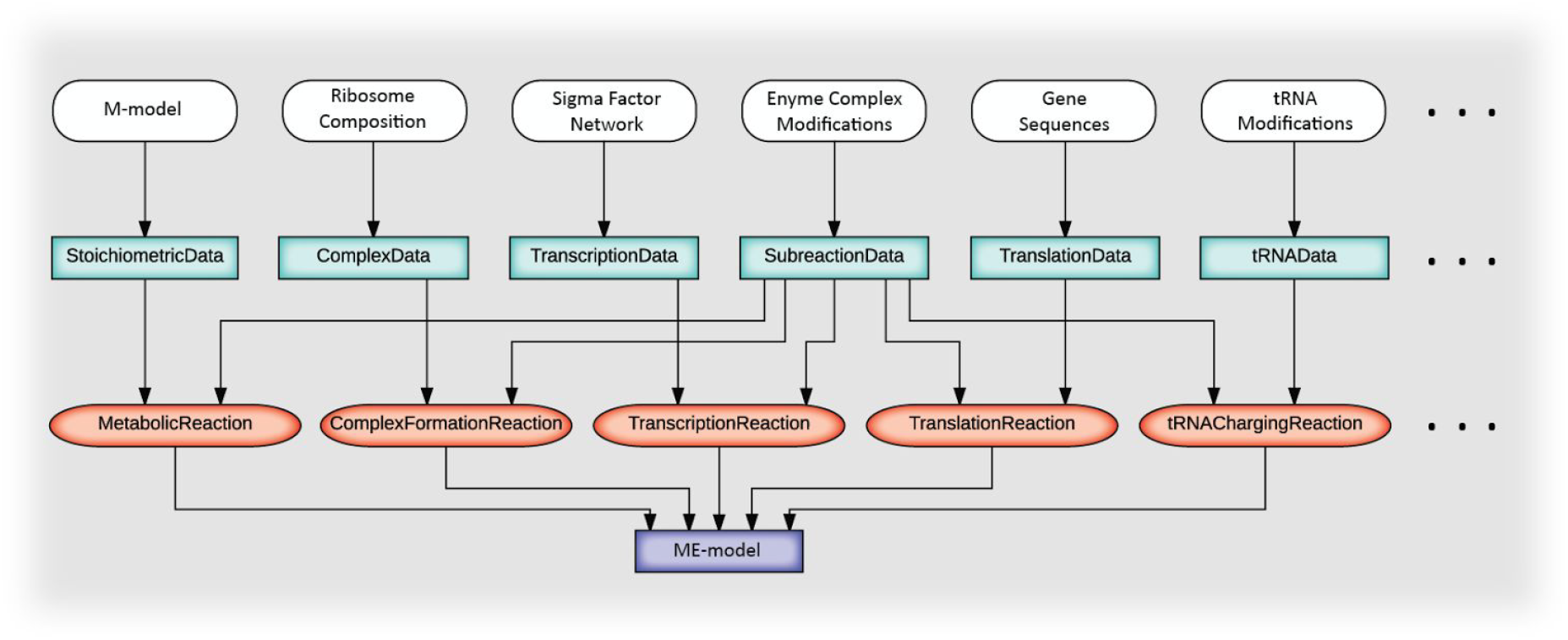
The flow of information from input data to the ME-model, as facilitated using the ‘**build_me_model**’ script. The ‘**build_me_model**’ workflow uses the ECOLIme package to load and process the *E. coli* M-model along with all supplied files containing information defining gene expression processes/reactions. This information is then used to populate the different ProcessData classes (shown in turquoise boxes) and link them to the appropriate MEReaction classes (shown in red ovals), all of which are defined in the COBRAme package. The entirety of the MEReactions comprise a working ME-model. Not all input data, ProcessData classes and MEReaction classes care shown. For a complete list, reference the COBRAme Documentation.

### Reformulating the *E. coli* ME-model

Significant efforts were made to simplify the ME-model while also optimizing the model size, modularity and time required to solve. These included: **1)** reformulating the implementation of explicit coupling constraints (metabolites) into single ME-model reactions and **2)** lumping major cellular processes such as transcription and translation into single ME-model reactions. Further, a number of updates, changes and corrections have been made to the *E. coli* ME-model reconstruction which are detailed below.

#### Macromolecular Coupling

The largest mathematical difference between the original ME-model formulation [6] and ME-models constructed using COBRAme is the change in the macromolecular coupling implementation. These coupling coefficients dictates the amount of macromolecule synthesis flux that is required for the reaction catalyzed by that macromolecule to carry flux. They are derived based on the fact that as a cell grows and divides it must dilute macromolecules its daughter cells and therefore have a general form of “μ/k_keff_” (O'Brien et al. 2013) (**Figure 3**). While these are essential in a ME-model to couple together the various reaction types, in previous model versions they inflated the number of metabolites and reactions contained in the ME-matrix, resulting in longer solve times. COBRAme improves coupling constraint implementation by directly embedding macromolecule dilution coupling into its catalytic reaction (**Figure 3**).

**Figure 3:**
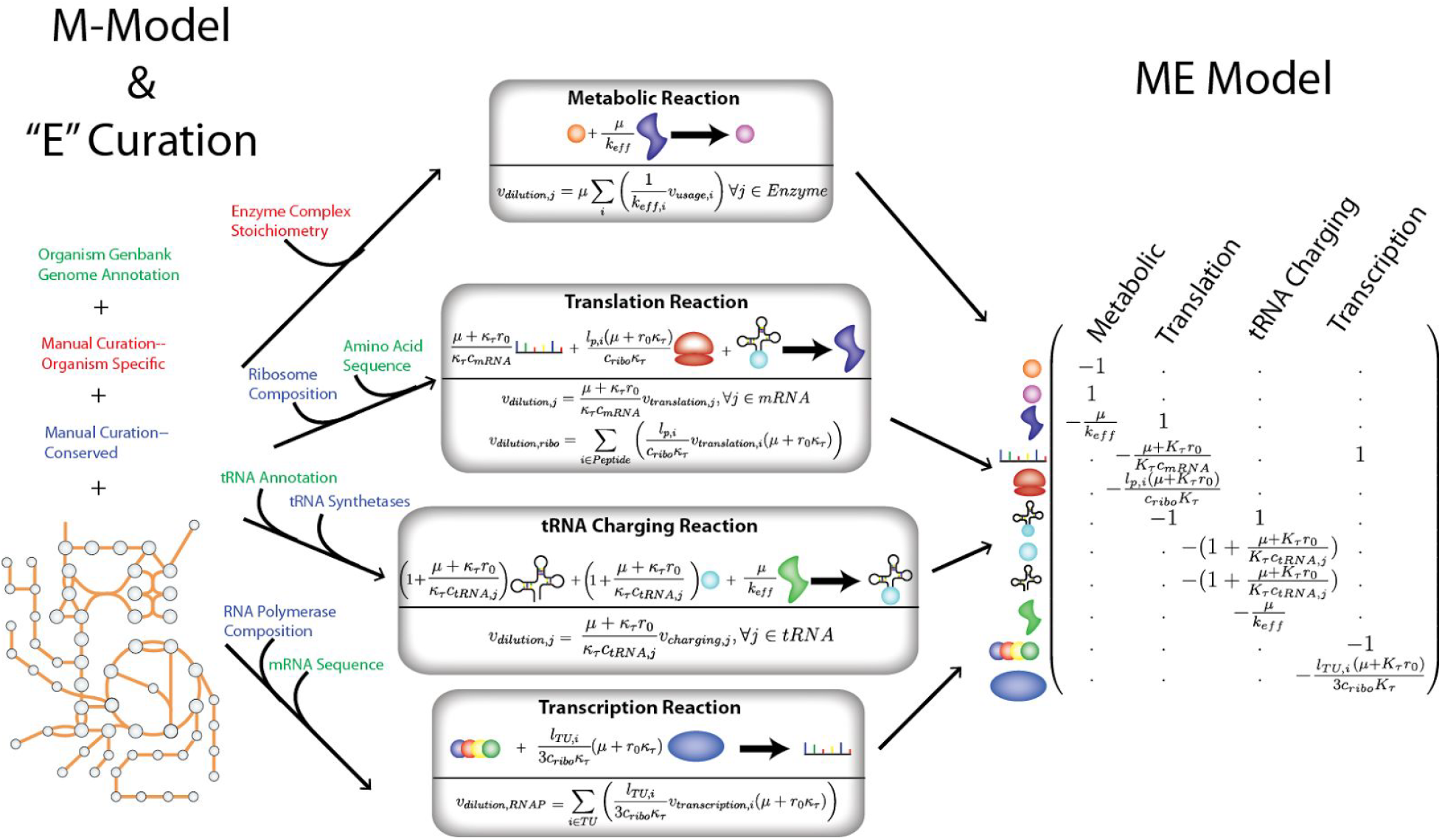
An overview of the COBRAme ME-model formulation. The previous ME-models implemented coupling constraints explicitly as model metabolites. With COBRAme, instead of using explicit coupling constraints (metabolites), dilution of coupled macromolecules to the daughter cell is accounted for by embedding them directly in the reaction which they are used. For example, for the metabolic reaction shown above, a small amount (μ/k_eff_)ofthe catalyzing enzyme is consumed by the reaction it is involved in. In other words, for each unit of flux carried by the metabolic reaction, μ/k_eff_ * v_metabolic_reaction_ of the catalyzing enzyme must be synthesized. A summary of the major macromolecular coupling that is accounted for in *i*JL1678b-ME is also shown, along with their representation in the ME-matrix.

The coupling coefficients were derived as in *O'Brien et. al. 2013*, but were effectively imposed in the ME-model as equality constraints (**Figure 3**) as opposed to inequality constraints used in previous ME-model implementations [6,7]. This means that each ME-model solution will synthesize the exact amount of each macromolecule as dictated by each coupling coefficient, thus giving the computed optimal macromolecule synthesis fluxes for the *in silico* conditions. Previous ME-model formulations, as stated above, have applied the constraints as inequalities thus allowing the simulation to synthesize macromolecule components above the value dictated by the coupling coefficient. While all enzymes are not fully saturated in *E. coli in vivo*, this phenomenon would not be selected as an optimal ME-model solution. Furthermore, using inequality constraints greatly expands the size of the possible solution space, significantly increasing the time required to solve the optimization. Reformulating the model using equality constraints thus resulted in a reduced ME-matrix with the coupling coefficients embedded directly into the reaction in which they are used (**Figure 3**). This process further removes the coupling constraints (metabolites) and associated variables (reactions) in the original formulation, which makes the ME-matrix much smaller (i.e. imposing each macromolecule coupling constraint in *i*JL1678b-ME requires at least one less variable (reaction) and constraint (metabolite) than previous ME-model versions).

Beyond reducing the size of the ME-matrix, eliminating inequality constraints greatly reduces the space of feasible fluxes at suboptimal growth rates. At an optimum, a ME-model will not waste resources, and, as a result, all the computed values are pushed up against their inequality constraints, rendering them as equalities. Therefore, reformulating the model with equalities alone will compute the same optimal flux state but results in a much simpler problem from a numerical point of view when applying the binary search or bisection solving algorithm (see **Optimization Procedure**).

To improve the modularity of the model the tRNA coupling was reformulated (see **S**). This was done in order to account for the dilution coupling of both the tRNA synthetase and tRNAs themselves during tRNA charging reactions. In doing so, the tRNA charging reactions now produce a charged tRNA equivalent that can be added to the translation reaction directly since dilution coupling has already been applied. This allows tRNAs to be cleanly removed without breaking the model by simply deleting the SubreactionData ProcessData instances that apply the tRNA equivalents.

A more thorough description of coupling constraints and their implementation can be found in the **COBRAme Documentation**.

#### Reaction Lumping

Using equality constraints in the COBRAme formulation and splitting the model into ProcessData and MEReactions allows for a variety of model simplifications. One major simplification is that reactions which occur in a number of individual steps or sub-reactions (i.e., ribosome formation, translation, etc.) can be lumped into a single reaction. The single MEReaction is constructed using the set of ProcessData instances that detail the individual sub-reactions involved in the overall reaction. This information is further accessible through the MEReaction instances itself which allows the information to be queried, edited and updated throughout the reaction. If the sub-reaction participates in many different reactions, the changes can be further be applied throughout the entire model. This lumping has the obvious benefit of reducing the number of model reaction, thus shortening the solve time. Lumping complex reactions has the added benefit of making the ME-model much more modular in nature. This simplifies the process of adding or removing new processes associated with the reaction. Examples of accessing and editing ProcessData through MEReactions can be found in the **COBRAme Documentation**.

#### Nonequivalent Changes

Unlike the reformulations described above, some of the changes made in the COBRAme formulation purposefully changed the model in a nonequivalent way. One of the most significant differences was assigning a “dummy complex” monomer with a representative amino acid composition as the catalytic enzyme for “orphan” reactions. These are non-spontaneous reactions which do not have a known enzymatic catalyst. The previous formulation therefore resulted in a slight bias toward using these reactions, given that they did not have an associated protein expression cost, which was corrected in *i*JL1678b-ME. Additionally, in *i*JL1678b-ME, protein “carriers” (e.g. acyl carrier protein) are assigned as catalysts to their transfer reactions. Therefore, the *i*JL1678b-ME will require translation of these carriers in order for them to participate in the reactions they are involved in, thus resulting in the expression of 52 more genes when simulating on glucose minimal media compared to *i*JL1678-ME.

Previous *E. coli* ME-models included individual reactions for all possible different combinations that a transcription unit could be excised of rRNA, tRNA or ncRNA. Though only 82 transcription reactions in are present in *i*JL1678b-ME, accounting for all excision possibilities and the subsequent degradation of the excised portions adds thousands of reactions to the model (a toy example is shown in the **Supplementary Text**). This has a sizeable effect on the solve time of the ME-model with limited improvement in the predictive capability of the model therefore handling this was removed from *i*JL1678b-ME.

Further, membrane surface area constraints imposed in *i*JL1678-ME were removed. This constraint limited the number of membrane proteins that could be expressed at a given growth rate. Protein competition for membrane space may play an important role in shaping *E. coli*’s metabolic phenotype, particularly when growing aerobically. Despite this, the constraint was removed to prevent the model from being over constrained when growing in non-glucose aerobic conditions, leading to unrealistic behavior. Removing this constraint makes *i*JL1678b-ME more generalizable. Similarly, growth-dependent surface area calculations were used when imposing lipid demands, therefore they were also removed and replaced with demands identical to those defined in the iJO1366 biomass objective function. The protein translocation genes and pathways added when reconstructing *i*JL1678-ME, however, remain in *i*JL1678b-ME.

Corrections were made to *i*JL1678-ME to remove metabolites from the *in silico* growth media that are not present in minimal *E. coli* growth media. For instance cob(I)alamin is not essential for growth in *E. coli*, but was required in the *in silico* growth media of *i*JL1678-ME to produce a feasible model. This was due to cob(I)alamin being an essential modification for the QueG complex. It has been shown that the presence of cob(I)alamin increases the epoxyqueuosine reductase tRNA modification activity, but is not required for the reaction to take place[24]. Further, a known failure mode of *i*OL1678-ME and *i*JL1678-ME is that all RNA modification genes are computationally essential, when, *in vivo*, this is not always the case. This means that the tRNA modification catalyzed by QueG is computationally essential. Given that the presence of cob(I)alamin in the *in silico* media will allow the activity of three reactions that would not be active when grown in minimal media (METS (catalyzed by MetH), ETHAAL, and MMM), the cob(I)alamin modification was removed from QueG and replaced with the two known iron sulfur cluster modifications. Biotin was also removed from the *in silico* growth media since *i*JL1678b-ME can synthesize it from glucose, therefore it does not need to be supplemented for growth.

### Optimization Procedure

Unlike M-models, the stoichiometric matrix for each ME-model consists of numerous growth rate (μ) dependant metabolite coupling coefficients and variable bounds (**Figures 1, 3**). This makes the ME-matrix nonlinear meaning they cannot be solved as a normal LP like M-models. The ME-matrix, however, is quasi-convex [25], meaning that, for any feasible substituted μ, all smaller μ values will also be feasible. Therefore, the maximal feasible μ value can be determined by a binary search or bisection algorithm wherein successive linear programs are solved at different values of μ to find the largest value of μ that gives a feasible flux state, as done for *i*JL1678-ME and *i*OL1650-ME. For each optimization the production of a representative dummy protein is maximized. In doing so, this allows the same algorithm to be used for both batch and nutrient limited growth, which required different procedures in *i*JL1678-ME and *i*OL1650-ME (O'Brien et al. 2013) (see **Supplementary Text**).

To perform the binary search, the following procedure was implemented in COBRAme. First, each symbolic coefficient or reaction bound was compiled into a function by SymPy [20]. Then, a linear program was created and passed into the linear programming solver, with all of these symbolic functions evaluated to an initial μ value. Afterwards, for each instance of the binary search in μ, values in the linear program were replaced by the new μ value, and the problem was resolved using the last feasible basis.

While any linear programming solver supported by COBRApy [18] could technically have been used, ME-models are very ill-scaled [6], unlike M-models. Therefore, two specialized solvers are used due to their extended numerical precision, thus ensuring acceptable numerical error: qMINOS[23,26] which supports quad (128-bit) numerical precision and SoPlex [22] which supports “long double” (80-bit) numerical precision as well as iterative refinement in rational arithmetic to further reduce numerical error.

## Results and Discussion

### Model Overview

The COBRAme framework was used to reconstruct a mass-balance checked, reformulated version of the *E. coli* K-12 MG1655 ME-model *i*JL1678-ME, called *i*JL1678b-ME. This produced a model with 12,654 reactions and 7,031 metabolites, a marked improvement over *i*JL1678-ME which contained 79,871 reactions and 70,751 metabolites. As a result, *i*JL1678b-ME has a matrix with ~90% fewer columns than *i*OL1650-ME. This dramatically speeds up the solving procedure and allows processes such as iterative refinement, which uses rational arithmetic and is unsuited for fast vector SIMD operations, to become feasible for fast and accurate solutions (**Figure 4**).

**Figure 4:**
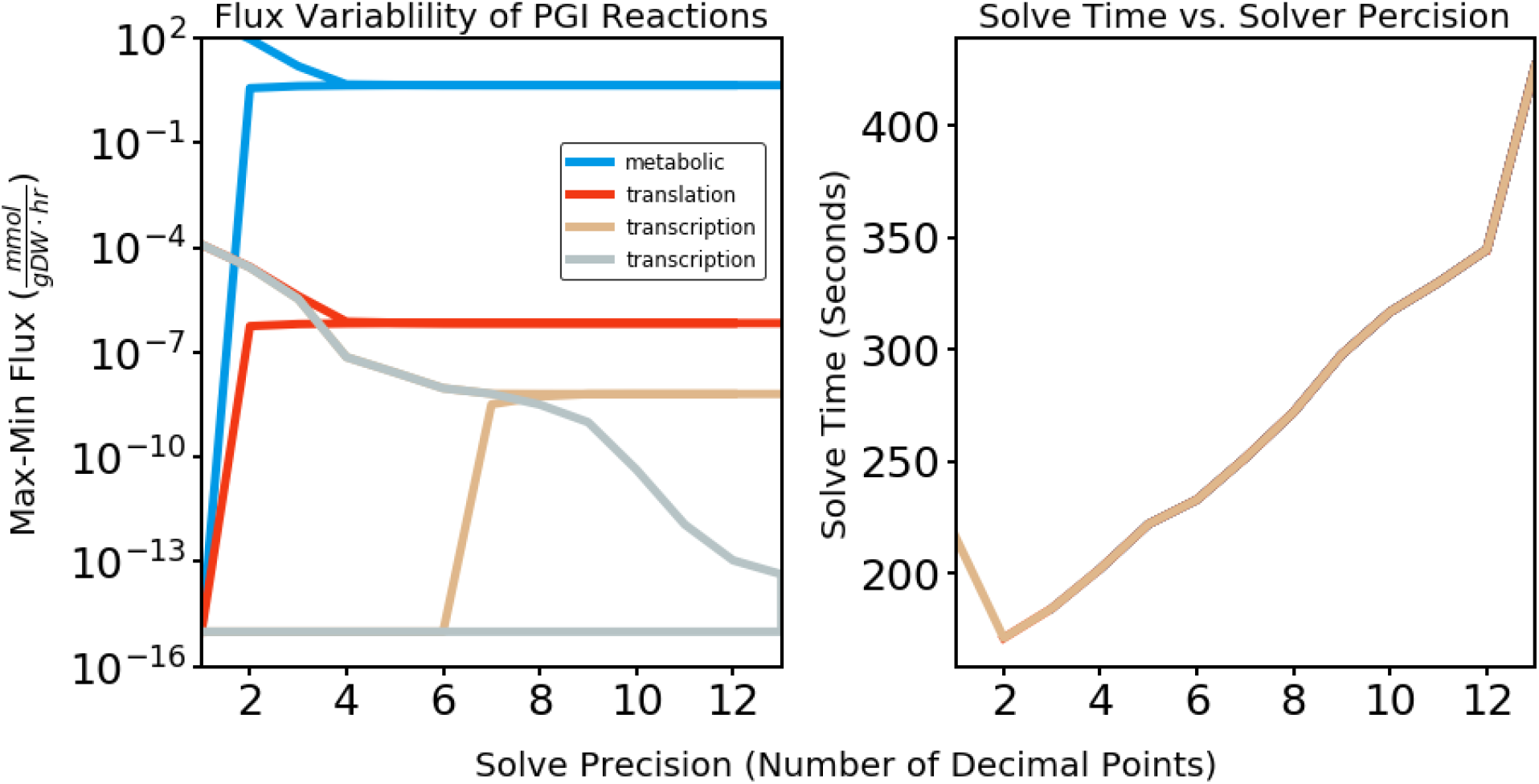
Flux variability analysis of reactions representing the expression of Pgi and the PGI metabolic reaction. With increasing solver precision, the variability becomes negligible (the max and min possible fluxes converge) for metabolic and translation fluxes when using a μ precision of 10^−5^ and for transcription fluxes when using a μ precision of at least 10^−10^. There are two transcription reactions for *pgi* to model transcription of this gene using two different sigma factors. High μ precision can be achieved without sizeable increases in total solve time, using qMINOS.

*i*OL1650-ME, constructed using COBRAme, was simulated in glucose aerobic minimal media *in silico* conditions and compared against simulations from the previous *i*OL1650-ME version. Both simulations were ran using a selection of k_eff_ parameters that were fit to proteomics data obtained from *E. coli* grown in multiple conditions [27]. The new model version gave very similar (R^2^>.98) fluxes when comparing model solutions on a transcription, translation and metabolic level (**Figure 5**) suggesting that the two models are virtually identical, computationally. The reformulated ME-model cannot be expected to give completely identical solutions as *i*OL1650-ME due to some of the nonequivalent changes and model corrections described in **Nonequivalent Changes**. Particularly, the RNA degradosome and RNA excision machinery was slightly under expressed due to the change in stable RNA excision handling described above.

**Figure 5:**
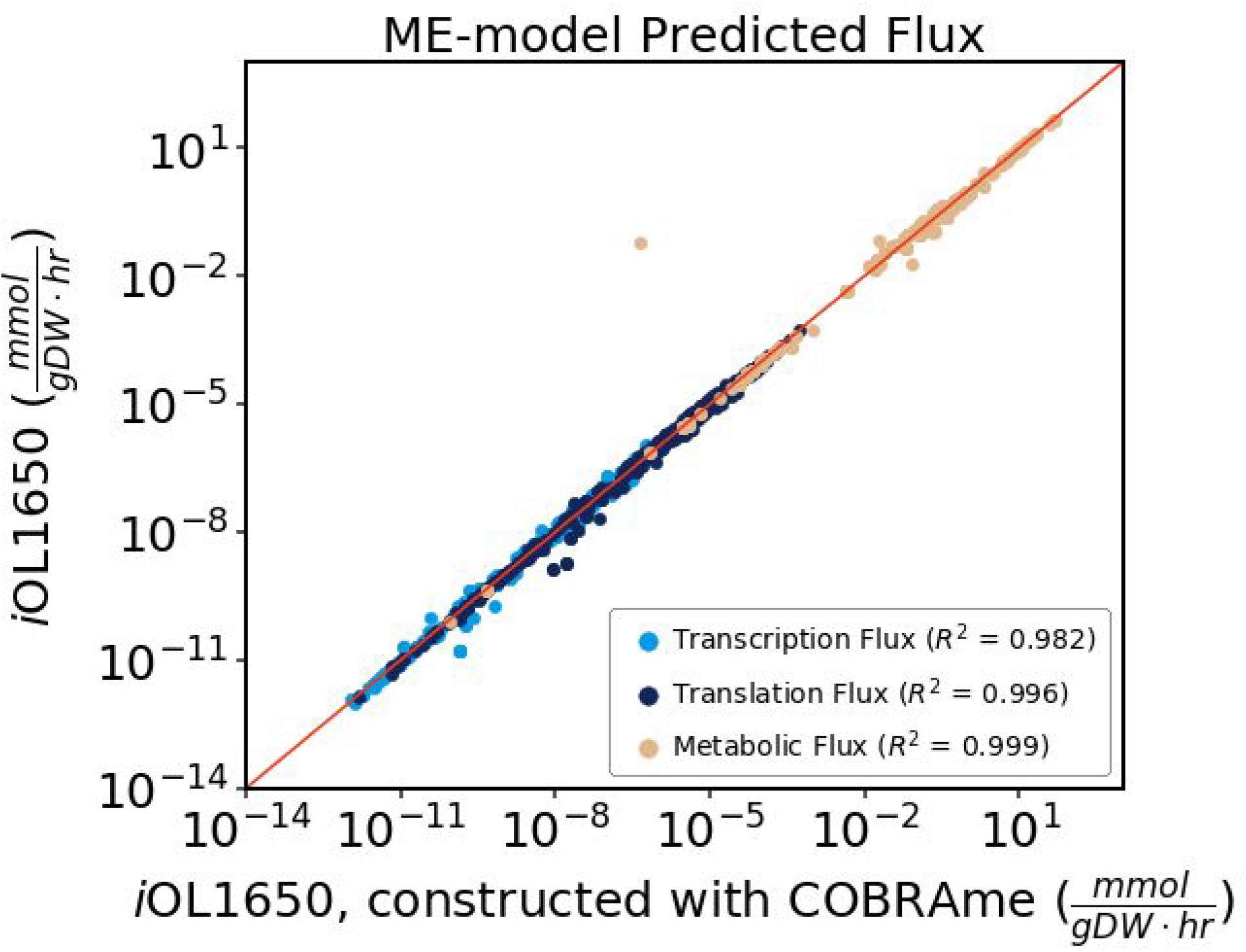
Comparison of the simulated fluxes of *i*OL1650-ME to the COBRAme generated version of the same model at transcription, translation and metabolic flux scales. At each level, the models provided comparable flux predictions, indicated by the Pearson Correlations above 0.98. The models cannot be expected to give completely identical flux predictions due to the ME-model updates outlined in **Nonequivalent Changes**. Since *i*JL1678b-ME does not contain membrane surface area constraints, *i*OL1650-ME was used for comparison.

Computational essentiality predictions for both *i*JL1678b-ME and *i*OL1650-ME were compared against a genome-wide essentiality screen of single gene knockouts grown in glucose M9 minimal media [28]. Due to the corrections described above, *i*JL1678b-ME displayed improved gene essentiality predictions when comparing essentiality for the 1539 proteins also modeled in *i*OL1650. The bulk of the these improvements stem from modeling the expression of enzyme “carriers” as mentioned in **Nonequivalent Changes.** This correction led to a 35 gene decrease in the number of false positives predictions made by *i*JL1678b-ME, but also led to a 22 gene increase in true positives. Overall the accuracy of the model improved from 86.6% to 87.5%. Further, the matthews correlation coefficient [29], a machine learning metric to gauge the performance of binary classifiers, saw an increase of 9.5% from 0.608 to 0.666.

Beyond performance and predictive capabilities, the reformulations and reduced size make *i*JL1678b-ME more understandable to the user. By lumping cellular processes into individual model reactions, the structure of the ME-model reactions is able to more closely resemble the central dogma of biology. For instance, the translation of a given gene, <gene_id> occurs in a single model reaction, “translation_<gene_id>” where all components and coupling constraints are applied in one place (**Figure 3**) as opposed to occurring in multiple separate reactions. In addition to being more easily understandable by the user, the reformulation makes the model more amenable to visualization tools like escher [19], further easing the process of interpreting simulation results.

### Editing a Constructed ME-model

The separation of COBRAme ME-models into information storing ProcessData classes and functional MEReactions allows many components of a finalized ME-model to be easily queried, edited and updated throughout the ME-model. This is especially useful for a few reasons: **1)** certain processes occur repeatedly throughout the process of expressing the genes and proteins within a cell. Therefore, if the user wants to edit a parameter associated with one of these processes, such as the GTP cost of translation elongations, this can be done by editing the ProcessData instance that defines this process and updating it through the model. **2)** Aspects of processes involved in gene expression can often be interrelated. For instance, the nucleotide sequence of an mRNA being translated dictates the number and type of charged tRNAs that are incorporate, the number of GTP driven elongation steps that must occur, etc. COBRAme allows the user to edit the sequence in one place and use the TranslationReaction’s update method to apply the changes throughout the reaction. Examples of making edits to a ME-model can be found in the **COBRAme Documentation**.

## Availability and Future Directions

Both the COBRAme and ECOLIme software packages are required to construct *i*JL1678b-ME and are currently available on the Systems Biology Research Group’s Github page (github.com/SBRG). Installation procedures as well as all necessary documentation required to build, simulate and edit ME-models are present in the repository READMEs. The SoPlex solver can be found at (http://soplex.zib.de/) and is freely available to all academic institutions. The soplex_cython package contains instructions to compile the soplex solver with 80-bit precision capabilities along with the necessary code required to solve *i*JL1678b-ME with SoPlex. Alternatively, the qMINOS solver[26] is also freely available for academic use. Instructions for installing and using the solver can be found as part of the solveme package[25]. These software packages will be actively maintained and improved. The COBRAme documentation can be found on readthedocs.

### Enable New ME-Model Reconstructions

We anticipate that the presented software tools will facilitate the reconstruction of many new ME-models beyond *i*JL1678b-ME for *Escherichia coli* K-12 MG1655. While the COBRAme code was constructed to be readily generalizable to many different organisms, it is likely that some organisms will require additional features for their ME-model reconstruction that we did not originally anticipate. It is our priority to continue to update and improve the code to enhance its utility to model new, diverse organisms. Future efforts will be also be made to create standards to govern how ME-models are reconstructed, structured and shared within the scientific community.

**Table 1:** Summary of essentiality predictions for the 1539 proteins modeled in both *i*JL1678b-ME and *i*OL1650-ME. Predictions of essentiality are from a genome wide screen of Keio collection[30] knockouts grown on glucose M9 minimal media [28].

|  |  | Experimental |  |
| --- | --- | --- | --- |
|  |  | Essential | Nonessential |
| ***i*JL1678b** | **Essential** | 1070 (69.5%) | 109 (7.1%) |
|  | **Nonessential** | 84 (5.5%) | 276 (17.9) |
| ***i*OL1650** | **Essential** | 1092 (71.0%) | 87 (5.7%) |
|  | **Nonessential** | 119 (7.7%) | 241 (15.4%) |

## Acknowledgements

We would like to thank Joshua Lerman, Aarash Bordbar, Justin Tan and Bin Du for informative discussions. This research used resources of the National Energy Research Scientific Computing Center, which is supported by the Office of Science of the US Department of Energy under Contract No. DE-AC02-05CH11231. We thank the Novo Nordisk Foundation [NNF16CC0021858] and the National Institute of General Medical Science of the National Institute of Health (award U01GM102098) for funding this project. CJL was supported by the National Science Foundation Graduate Research Fellowship under Grant no. DGE-1144086

